# Single-cell RNA-seq reveals identity and heterogeneity of malignant osteoblast cells and TME in osteosarcoma

**DOI:** 10.1101/2020.04.16.044370

**Authors:** Yan Zhou, Dong Yang, Qing-Cheng Yang, Xiao-Bin Lv, Wen-Tao Huang, Zhenhua Zhou, Ya-Ling Wang, Zhichang Zhang, Ting Yuan, Xiaomin Ding, Li-Na Tang, Jian-Jun Zhang, Jun-Yi Yin, Yu-Jing Huang, Wen-Xi Yu, Yong-Gang Wang, Chen-Liang Zhou, Yang Su, Ai-Na He, Yuan-Jue Sun, Zan Shen, Bin-Zhi Qian, Peizhan Chen, Xinghua Pan, Yang Yao, Hai-Yan Hu

## Abstract

Osteosarcoma (OS) has high heterogeneity and poor prognosis. In order to explore the molecular mechanism of OS and the tumor micro-environment (TME) on OS, we employed single-cell RNA-sequencing (scRNA-seq) on 110,745 individual cells from OS primary lesion, recurrent focal and metastatic tissues. We identified 5 main malignant subpopulations of OS cells, 3 clusters of osteoclast(OC) and 2 types of cancer-associated fibroblasts (CAFs). Further we found that the progenitor OC and, antigen presenting CAF (apCAF) were lower in lung metastatic and recurrent tumor tissues than in primary tumor tissue. M2-like macrophages were predominant in the TME myeloid cells. Inactivation state of tumor-infiltrating T cells, mainly the CD4-/CD8-T and Treg cells, existed in lung metastatic tissues. T-cell immunoreceptor with Ig and ITIM domains (TIGIT) expressed in 11 samples. We then blocked TIGIT which significantly enhancd the cytotoxic effects of primary T cells on OS cell lines. Our report represents the first use of scRNA-seq for the transcriptomic profiling of OS cells. Thus, the findings in this study will serve as a valuable resource for deciphering the intra-tumoral heterogeneity in OS and provide potential therapeutic strategies for OS in clinic.

## Introduction

Osteosarcoma (OS) is a highly aggressive malignant bone tumor frequently occurring in children and adolescents[1–3]. The incidence of OS is about 4.8 per million per year. Traditionally, the standard treatment protocol for OS consists of extensive surgical resection, chemotherapy, and radiation. Researchers have been screening the effective target drugs on OS for decades. In recent years, the vascular endothelial growth factor receptor-tyrosine kinase inhibitors (VEGFR-TKIs) have appeared outstanding due to their effectiveness. However, as stated by the Surveillance, Epidemiology, and End Results (SEER) Program, the five-year overall survival rate for patients with bone sarcoma is 66.2% (2009 to 2015) [4]. According to the published data, the relapse and/or metastasis rate of OS remains to be higher than 30%. For these patients, the five-years overall survival rate was even worse, being about 10–30% [5]. As such, there is an urgent need to identify the molecular mechanism and novel therapeutics that may improve management of OS.

Immune checkpoint inhibitors have led to a breakthrough in immunotherapy for a variety of solid tumors [6, 7]. However programmed cell death 1 (PD-1) inhibition has limited effect in OS[8, 9]. Davoli revealed that highly aneuploid tumors showed reduced expression of markers of cytotoxic infiltrating immune cells, especially CD8+ T cells, and increased expression of cell proliferation markers. Immune evasion markers correlated mainly with arm- and chromosome-level somatic copy number alterations (SCNAs), consistent with a mechanism related to general gene dosage imbalance rather than the action of specific genes. In this regard, OS is the typical one [10]. Previous reports show that OS is abundant in widespread and recurrent somatic chromosomal lesions, including structural variations (SVs) and copy number alterations (CNAs); however, few recurrent point mutations in protein-encoding genes have been identified in OS[11–15].Low expression of immune-associated genes is another significant phenotype for OS[16]. How to convert the immunosuppressive microenvironment into the one that favors the induction of antitumor immunity is indispensable for effective cancer immunotherapy.

Here, we employed single cell transcriptome approach to dissect the heterogeneity of OS cells. We analyzed the transcriptomic profiles of a total of 110745 cells from 7 primary tumors, 2 lung metastatic and 2 recurrent OS tissues. We first divided the OS cells into 5 sub-clusters and osteoclast into 3 subtypes. The profiles of OS, OC and immune-system cells were analyzed. We found that the TME of the recurrent and lung metastatic OS exhibited more significant suppressivity than primary tumor tissue. Thus, the results in this study improve the understanding of the immune-suppressivity observed in advanced OS including distant metastasis and recurrence, and are potentially valuable in novel treatment strategy for OS.

Of importance, we are the first to uncover that Treg cells in OS expressed TIGIT. TIGIT is a coinhibitory receptor expressed on effector T cells, natural killer (NK) cells, T regulatory cells (Treg) and T follicular helper (TFH) cells. It has gained attention as a potential therapeutic target in wide variety of tumors [17–19]. The antibodies of TIGIT, named BGB-A1217, had registered and recruited on August 2019. Here we explored the preclinical significance of blocking of TIGIT.

## Method

### Patients

The eleven patients for scRNA-seq analysis enrolled in this study were hospitalized during the period of October 2017 to April 2019 in Shanghai Sixth People’s Hospital. The study was approved by Shanghai Sixth People’s Hospital Ethics Committee. Each patient was provided a written signed consent. All patients were diagnosed according to the National Comprehensive Cancer Network (NCCN) Clinical Practice Guidelines in Oncology with the terms of Bone Cancer (Version 2.2019). Among 11 patients for scRNA-seq, 7 were derived from the primary sites of patients who received traditional first line combination chemotherapy for OS, including Adriamycin(ADM), cisplatin(DDP), methotrexate(MTX) and Ifosfamide(IFO) and surgical therapy. 2 lung metastatic patients and 2 recurrent patients all received the gemcitabine combined with Docetaxel(GT) chemotherapy treatment. The BC17, one of the lung metastasis patients, had enrolled in clinical trial NCT03676985 and undergone the anti-PD-L1 treatment for 6 times. For this clinical trial, all enrolled patients had finished the neoadjuvant chemotherapy, operation and adjuvant chemotherapy. They would accept anti-PDL-1 for one year until the disease progresses. Detailed information of the 11 patients was provided in Table 1 respectively. Tow patient agreed to donate blood for us to explore the effect of anti-TIGIT.

**Table 1.**
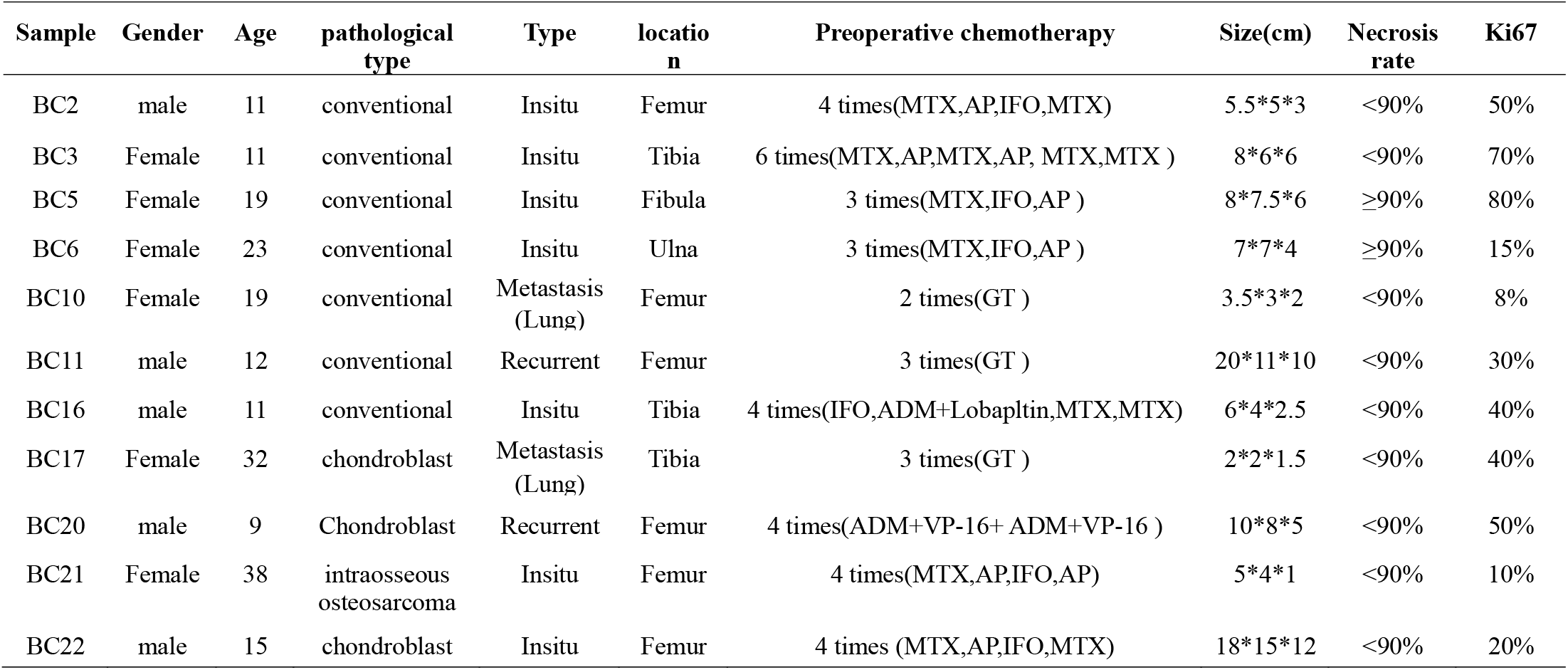
Clinical Characteristics of OS Patients

### Sample preparation and cell purification for scRNA-seq

The fresh tumor tissue was stored in the GEXSCOPETM Tissue Preservation Solution (Singleron) and transported to the Singleron lab on ice as soon as possible. The specimens were washed with Hanks Balanced Salt Solution (HBSS) for 3 times and minced into 1–2 mm pieces. Then the tissue pieces were digested with 2 ml GEXSCOPETM Tissue Dissociation Solution (Singleron) at 37°C for 15 min in 15 ml centrifuge tube with sustained agitation. After digestion, the samples were filtered through 40-μm sterile strainers and subsequently centrifuged at 1,000 rpm for 5 minutes. Thereafter, the supernatants were discarded, and the cell pellets were suspended in 1 ml PBS (HyClone). To remove the red blood cells, 2 mL GEXSCOPETM red blood cell lysis bu□er (Singleron) was added at 25°C for 10 minutes. The solution was then centrifuged at 500 × g for 5 min and suspended in PBS. The sample was stained with trypan blue (Sigma) and evaluated microscopically.

### 10x library preparation and sequencing

Single-cell suspensions were converted to barcoded scRNA-seq libraries by using the Chromium Single Cell 3’Library, Gel Bead & Multiplex Kit (10x Genomics, V3), and following the manufacturer’s instructions. Briefly, cells were partitioned into Gel Beads in Emulsion in the ChromiumTM Controller instrument where cell lysis and barcoded reverse transcription of RNA occurred. Libraries were prepared using 10x Genomics Library Kits and sequenced on Illumina HiSeq X with 150 bp paired end reads.

Raw reads were processed to generate gene expression profiles using an internal pipeline. Briefly, after filtering read one without poly T tails, cell barcode and UMI was extracted. Adapters and poly A tails were trimmed (fastp V1) before aligning read two to GRCh38 with ensemble version 92 gene annotation (fastp 2.5.3 a and featureCounts 1.6.2). Reads with the same cell barcode, UMI and gene were grouped together to calculate the number of UMIs per gene per cell. The UMI count tables of each cellular barcode were used for further analysis. Cell type identification and clustering analysis using Seurat program. The Seurat program (http://satijalab.org/seurat/, R package, v.3.0.1) was applied for analysis of RNA-Sequencing data.

### Cell-type identification and subgroup categorization by the t-SNE method

The individual Seurat objects were integrated with the SCTransformation algorithm provided by the Seurat package. The top 3,000 Highly variable genes across the cells were chosen to perform the Principal Component Analysis(PCA) analysis. Top 50 significant PCAs were applied for the graph–based clustering based on the t-SNE method to identify the main cell groups for all samples. In the subgroup cell identification, the top 10 PCAs were applied for the graph based clustering. The cells in each integrated subcluster were selected to run the SCTransformation analysis with the cells being categorized with the top ranked, differentially expressed genes and the well-known cellular biomarkers. The osteoclast cell biomarkers were defined as the cathepsin K5(CTSK5) and tartrate-resistant acid phosphatase(TRAP/ACP5); cadherin11(CDH11), Integrin Binding Sialoprotein (IBSP) for osteoblast cells; lumican(LUM), decorin(DCN), collagen, type I, alpha 1(COL1A1) for fibroblast; CD74, CD68 for monocytes; CD2, natural killer cell granule protein 7(NKG7) and CD3D for T and NK cells.

### Differential expressed genes(DEG) identification and Gene Ontology(GO) enrichment analysis

The cluster subgroup specific biomarkers were identified with the FindAllmarkers functions implemented in the Seurat package based on the normalized gene expression data. The genes with the adjusted P-values < 0.05 between the clusters were defined as DEGs and were selected for the GO enrichment analysis using the ClusterProfiler package of R.

### CNV estimation in the OS tumor

Initial CNVs for each region were estimated by inferCNV R package. The CNVs of total cell types were calculated by expression level from single-cell sequencing data for each cell with –cutoff 0.1 and –noise_filter 0.2. For each sample, gene expression of cells was re-standardized and values were limited as −1 to 1.

### Construction of single cell trajectories of OS cells

To identify genes that were involved in the progression of OS cells, the Monocle2 package (v2.8.0) was used to analyze single cell trajectories from primary tumor to the lung metastasis or recurrence. We used top 100 differentially expressed genes across the cell types identified by the Monocle 2 to sort the cells in pseudo-time order. The gene expression files in the primary cells were defined as the root_stage and the DDRTree was applied to reduce the dimensions and visualize the plot_cell_trajectory functions implemented in the monocle2. Differentially expressed genes over the Pseudo-time from primary tumor to lung metastasis or recurrent were calculated by the “differentialGeneTest” function in Monocle2 (q value < 10^−20^). The genes were categorized into 6 subgroups and the GO function enrichment analysis was performed for the genes in each cluster with the ClusterProfile package of R.

### Immunohistochemistry(IHC) staining and immunofluorescence(IF) staining

Tissue sectioning and IHC staining of formalin fixed paraffin-embedded (FFPE) OS specimens were performed following the general methods. All sections were deparaffinized, rehydrated, and washed and endogenous peroxidase was blocked using3% H_2_O_2_ for 10 min, the slides were incubated with primary antibodies followed by HRP-linked secondary antibodies and diaminobenzidine (DAB; ZhongShan Golden bridge biotechnology Co LTD, Cat No. ZLI-9018) staining. Counterstaining was done with hematoxylin. Slides were dehydrated with sequential ethanol washes for 1 min each starting with 75%, followed by 80% and finishing with a 100% ethanol wash. Two physicians blinded for clinical/tumor-characteristics independently assessed IHC-staining for TIGIT, CD3, CD4, CD8, CD74 and CTSK.

For IF staining, the process was same to above until inncubating with primary antibodies overnight at 4°C. Fluorescence-labeling secondary antibodies including donkey anti-rabbit Alexa Fluor488 (Molecular Probes, catalog A21202, 1:1000) and goat anti-mouse Alexa Fluor 514nm (Molecular Probes, catalog A31555, 1:1000) were incubated for 1 hour at room temperature after washing. Nuclei were counterstained with DAPI (MilliporeSigma, D9542). Sections were mounted using fluorescence mounting medium (Dako, S3023).

### Cytotoxicity assays by CytoTox 96^®^ Non-Radioactive Cytotoxicity Assay

PBMCs were collected from BC3 and BC16 by density centrifugation using Lymphocyte Separation Medium (MP Biomedicals). Then CD3+ T cells were isolated using the MACS positive selection technology (Miltenyi Biotec) according to the manufacturer’s protocol. For T-cell activation assays, CD3+ cells were seeded in 24-well plates and stimulated with IFN-y(1000 U/mL; Peprotech), IL-2 (600 U/mL, Peprotech) and anti-CD3 antibody (5 ng/mL, clone OKT3; Biolegend) for 3 days then blocked TIGIT with TIGIT antibodies(50 μg/ml, clone #A15153G, Biolegend) for 24h. 143B and U2OS cells were seed in 96-well plates overnight, then added CD3+ T cell at effector-to-target (E:T) ratios of 4:1 and 8:1. Co-culture system were incubated for 8 h. The supernatant was harvested and was subjected to analysis by the CytoTox 96^®^ Non-Radioactive Cytotoxicity Assay. The killing effect of T cells against target cells was assessed with the following equation: Cytotoxicity = (Experimental – Effector Spontaneous – Target Spontaneous)/(Target Maximum–Target Spontaneous) × 100%. All experiments were performed at least three times.

### Statistical analysis

Statistical analysis was performed using statistics package for social science 21.0 (SPSS 21.0; SPSS Inc, Chicago, IL). All the data were expressed as mean±SD. The significance was determined by the t test. *p*<0.05 was considered statistically significant.

## Results

### Single-cell analysis uncovers the complexity of OS tumor

To explore the cellular compositions in OS, we performed scRNA-seq analysis of 7 primary OS tumors, plus 2 recurrent OS tumors, and 2 samples from pulmonary metastasis (Table 1). After initial quality control, we acquired single-cell transcriptomes in a total of 110,745 cells, including 72,004 cells from *in situ* samples, 19,439 cells from lung metastasis samples, and 19,302 cells from recurrence samples. We first applied principle component analysis on variably expressed genes across all cells and identified six main cellular clusters including OS (osteoblast cell, 47,598), osteoclast cell (9,180), fibroblast (26,772), myeloid cell (18,158), endothelial (3,621) and T/NK cell (5,416) based on the t-distributed stochastic neighbor embedding (t-SNE) analyses in two dimensions (Fig. 1A). The t-SNE results for individual patients were shown in Supplementary Fig. S1. We performed the differential expression analysis to identify the cellular cluster-specific genes and defined the cellular cluster together with the well-known cellular biomarkers. The dot-plot and violin-plot showed the well-known cell type-specific markers embedded in cells from distinct clusters (Fig. 1B and C). The heatmap gathered the well-known cell type markers to distinguish each cell cluster, such as CTSK, ACP5 for osteoclast, CDH11 and BSP for malignant osteoblast, which we defined here as OS cell.

**Fig. 1.**
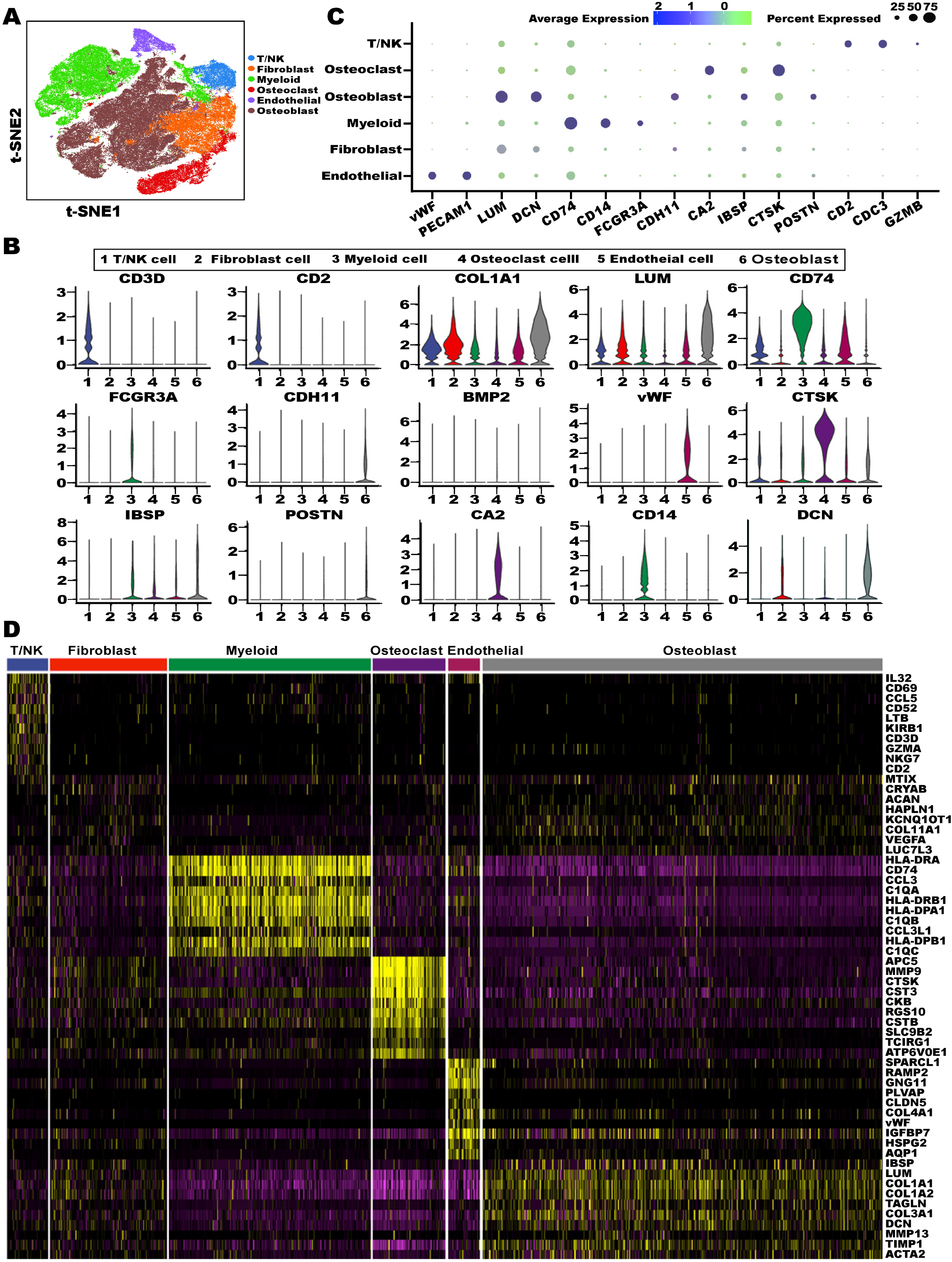
Delineation of diverse cell types in OS using the scRNA-seq method. (A) The t-SNE plot displayed the main cell types in OS tissue. (B) Violin plots demonstrated the expression levels of cluster-specific marker genes. (C) Dot plot displayed multiple well-known cell type-specific biomarkers across clusters. The size of dots represents the proportion of cells expressing a particular marker, and the spectrum of color indicates the mean level of this gene. Legends are shown as above. (D) Heatmap showed the significant gene in each cell group.

### Intra-tumoral heterogeneity in malignant OS cells

With the t-SNE analysis of osteoblast OS cells, we identified 5 distinct subgroups, named as metabolic, proliferating, extracellular matrix remodeling, ossification and cellular differentiation cells, respectively (Fig. 2A). Based on the GO analysis, cluster 1 was enriched in genes related to structural constituent of ribosome, glycolysis and active lipid metabolism and therefor termed as metabolic OS cells. Cluster 2 was enriched in genes related to cell cycle and proliferating with relatively higher expression of Ki67 and TOP2A and termed as proliferating OS cells. Cluster 3 was enriched with genes related to extracellular matrix modeling and therefor defined as ECM modeling OS cells. Genes in cluster 4 had a high level of genes involved in sulfur compound binding, heparin binding and glycosaminoglycan binding pathway, suggesting it was associated with ossification and therefor defined as ossification. Cells in cluster 5 were specific in cell differentiation processes including histone acetyltransferase binding, RNA polymerase II transcription factor binding and DNA-binding transcription activator activity. Many transcription factors, such as JUN, MYC, SOX9, etc, were over-expressed, suggesting that they may be pluripotent (Fig. 2B), and the cells were defined as cellular differential OS cells. The KEGG analysis showed that the genes of TP53 pathway were markedly disordered in subgroup 5 (Fig. S2A). The heatmap displayed the key genes characterizing our classification (Fig. S2 B).

**Fig. 2.**
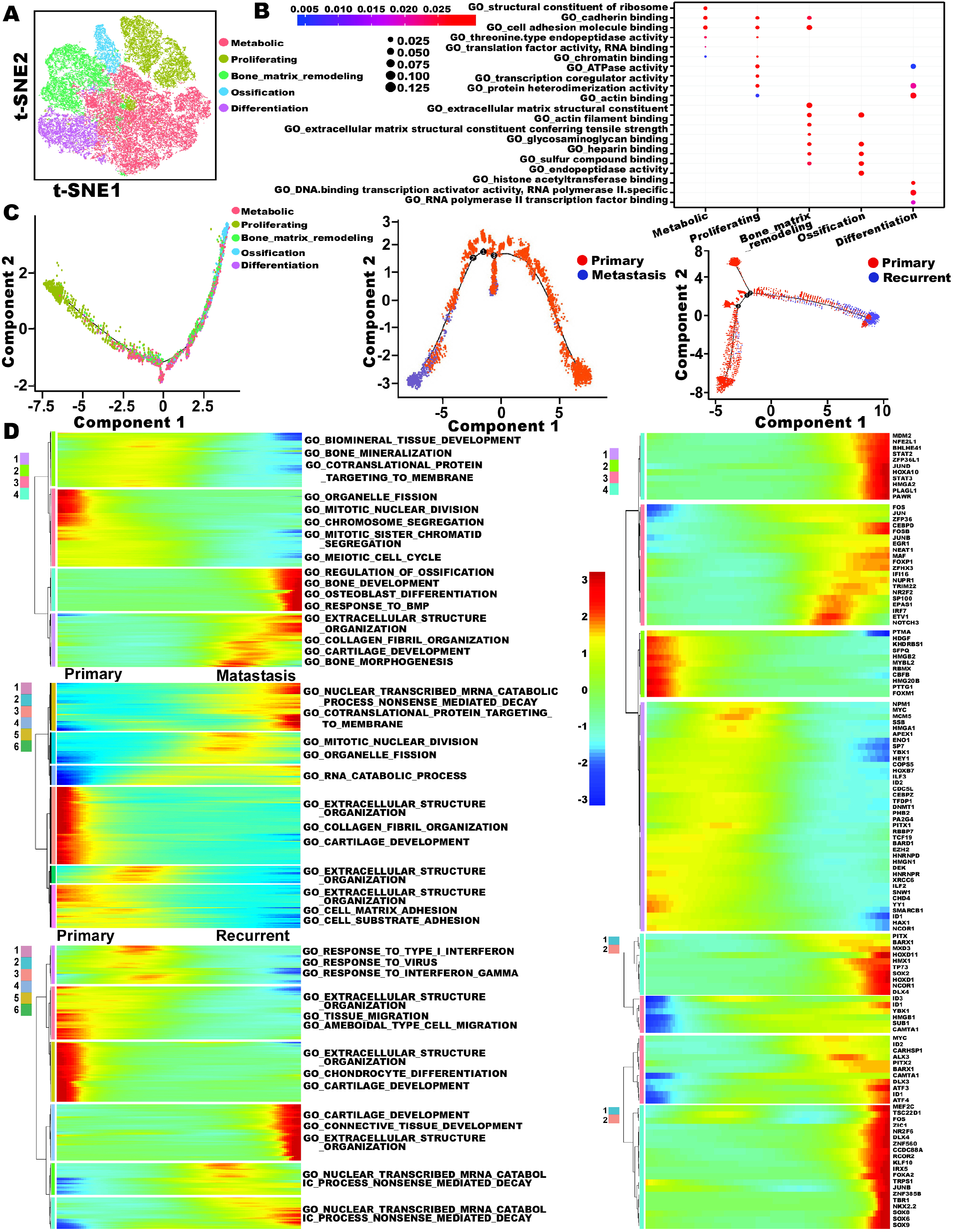
Differential gene expression profiles in malignant OS cells. (A) The malignant osteoblast cells, here named OS, were divided into 5 subtypes based on t-SNE analysis. (B) The characteristics of 5 subgroups. OS cells were differentiated by GO analysis. (C) The trajectory analysis of the OS cells included primary, metastasis and recurrence. (D) The primary OS cells gathered 4 clusters. For primary and metastatic OS cells, the differentially expressed genes (rows) in conformity with the pseudo-time (columns) gathered hierarchically into 6 cluster profiles. For primary and recurrent OS gene profiles, 6 clusters were gathered. Furthermore, we outlined the corresponding diagram on the basis of the transcription factor data. Color key from blue to red indicates relative expression levels from low to high.

To address the origin differentiation, development and stemness of OS, we performed the trajectory analysis of OS cells(Fig. 2D). Firstly we evaluated the genes that expressed along with the pseudo-time in primary OS cells. It is uncertain which cell type is responsible for OS initiation. Our data imply that in the primary OS tissue, the highly proliferating OS cells would transit to the differentiation cells with transcription factor including, over-expression and then transform to special potential cells including hypermetabolism, bone matrix remodeling and ossification. We further analyzed the gene patterns along with the cellular trajectories, and the genes were subclustered into 4 groups. The GO analysis suggested that the genes related to mitotic nuclear division were down-regulated along with the trajectory while genes related to regulation of ossification and bone morphogenesis were increased along with the trajectory.

Secondly, we performed the cellular trajectories from primary to lung metastasis. In the lung metastasis, the genes were categorized into 6 clusters, with the genes related to cellular matrix being significantly down-regulated, while the genes related to mitotic nuclear division, organelle fission, RNA catabolic process, nuclear transcribed mRNA catabolic process nonsense mediated decay and cotranslational protein targeting to memberane *etc*. were significantly increased. Furthermore, ten transcriptional factors, including (SRY-like HMG box) SOX2, TP73, and homeobox gene family D11 (HOXD11) *etc*., were significantly increased in the lung metastasized cells, suggesting that they play important roles in lung metastasis of OS cells.

Thirdly, we explored the cellular trajectories from primary to recurrent OS cells. The genes in response to IFNγ were reduced while genes related to the connective tissue were significantly increased. Thirty-one transcriptional factors, including MYC, FOS, ORF of iroquois homeobox 1 (IRX5) and JUNB *etc*., were significantly increased in the local tumor recurrence, suggesting that they play vital roles in the diseases recurrence.

We also calculated large-scale chromosomal CNV in each subject based on averaged expression patterns across intervals of the genome (Fig. S3). We found that, while the genomic region of 8q was frequently increased in the OS cells, 6p region was frequently down-regulated, which were in consistent with previous studies performed with the CGH methods[20].

### From antigen presenting to bone resorption during the OC maturation

The bone or bone-like microenvironment/niches provide growth and survival signals essential for OS initiation and progression. OC is an important type of cells to maintain the balance of bone formation. Previous studies suggested that OC cells express immune regulators, uptake soluble antigens and secrete cytokine to activate both CD4+ and CD8+ T cells in an MHC-restricted fashion[21]. Mature osteoclast (OC) is a type of poly-nuclear cell involved in the bone resorbing. We hypothesize whether the function of OCs dynamic change with development. Here we divided the OC cells into 3 main subgroups based on the integration data, including progenitor OC, immature OC and mature OC cells(Figure 3A) The progenitor OC cells showed relatively higher CD74 and the topoisomerase IIα (TOP2A), while the mature OC cells displayed higher expression of CTSK and ACP5 (Fig. 3B). The cellular trajectory analysis is consistent with our hypothesis that the cells with higher CD74 expression level were ranked at the origin of the pseudo-time linkage while the expression of CTSK and ACP5 were increased along with the pseudo-time(Fig. 3C).

**Fig. 3.**
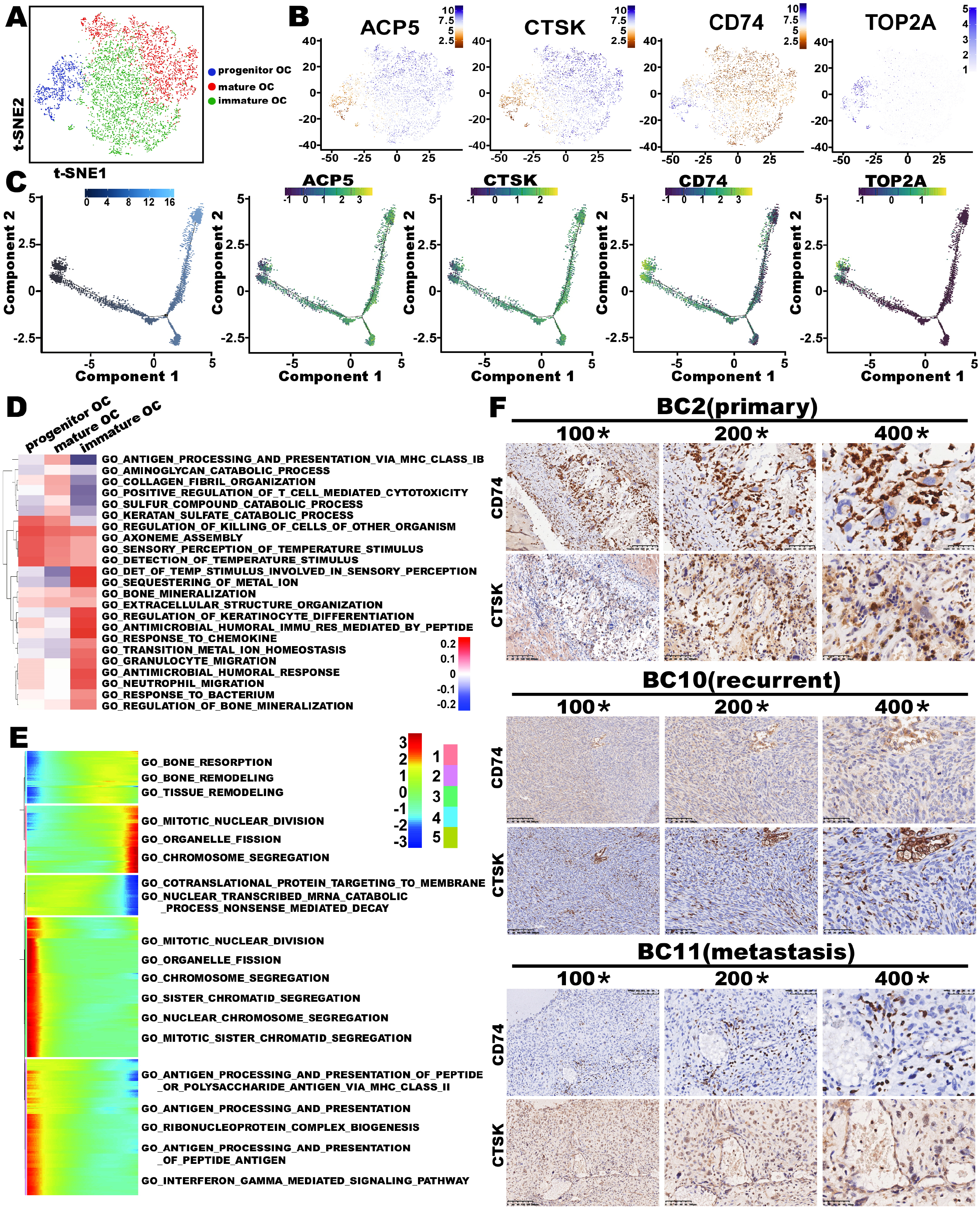
Distinct subpopulations of osteoclastic cells based on scRNA-seq data. (A) Graphical (t-SNE) plot demonstrated 3 main cell subtypes of OC. (B)We also showed the t-SNE plot figure marked ACP5, CTSK, CD74 and TOP2A separately. (C) Pseudo-time figure showed the development of OC subpopulations. We also presented the Pseudo-time marked ACP5, CTSK, CD74 and TOP2A respectively. (D) Different cell subtype clusters are color coded. The OC subtypes were classified according to the expression levels of specific genes represented in the heatmap. The GO analysis displayed the significant differences among 3 clusters. (E) The row graph was gathered hierarchically into 5 clusters. (F) We detected the expression of CD74 and CTSK in OS samples by IHC.

The primary function of CD74 is regulation of T-cell and B-cell developments, dendritic cell (DC) motility, macrophage inflammation, and thymic selection[22]. In addition CD74 can act as a receptor for macrophage migration inhibitory factor (MIF). It was found that MIF inhibited osteoclast formation and CD74 knockout (KO) mice had decreased bone mass[23]. We detected the CD74 and CTSK co-expression in OCs by IHC method on serial section. We found the cells with CD74 extreme positive were small and mononuclear with weakly positive of CTSK. The CD74 level in multinuclear OCs presenting light brown was markedly lower than in mononuclear OCs(Fig. 3D). The IF results were equal to IHC(Fig. S5). This result also indicated that the antigen presentation function of OC fade away with its development.

Furthermore, we analyzed the gene change in 25 GO biological process categories. The genes related to antigen processing and presentation via MHC class IB, aminoglycan catabolic process, collagen fibril organization *etc*, showed a significant increase along with the differentiation of OC cells, while the genes related to sequestering of metal ion, bone mineralization, extra cellular structure organization *etc*, were down-regulated with the differentiation (Fig. 3E, Fig. 3F). For our opinion, along with the differentiation of OC, the antigen presenting function was diminishing, while the bone resorption function became stronger and stronger. The deficiency of progenitor OC in metastasis and recurrent OS tissue may contribute to the immunosuppressed state.

### Distincted capCAFs in OS

CAFs modulate tumor stiffness and facilitate cancer progression. Based on the reported CAF biomarkers including land use Lumican(LUM), collagen type I alpha 1 chain(COL1A1) and decorin(DCN), a total of 18,158 fibroblast cells were identified. These fibroblast cells were categorized into two distinct subclusters (Fig. 4A), including myofibroblastic CAFs (myCAFs) with periglandular FAP+ αSMA^high^ and apCAFs with high level of MHC class II family members [24]. Compared to the myCAFs, the apCAFs showed relative higher expression level of CD74 and the MHC II molecules while the expression level of DCN and LUM was relatively lower (Fig. 4B). We then generated the heatmap according to cluster-specific marker genes by performing differential gene expression analysis to define the identity of each cell cluster (Fig. 4C). It’s worth noting the apCAFs were frequently identified in the primary tissue (1335/4587), but rarely noted in the metastasis (226/5000) and the recurrent tumors (452/5217), implying the antigen presenting was more active in primary OS tissue.

**Fig. 4.**
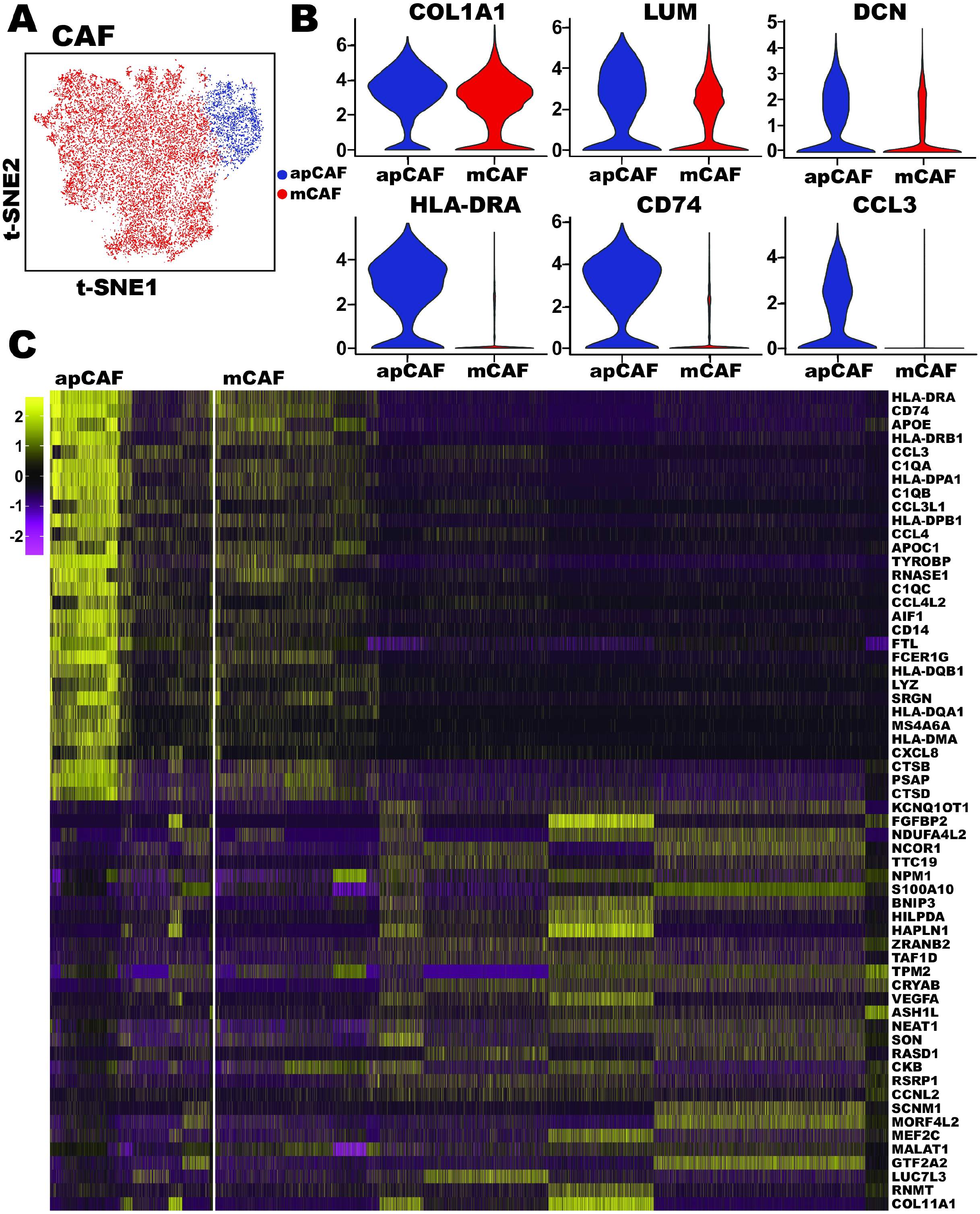
Identificaion of CAF subtypes in OS tissue. (A) Unsupervised clustering of two CAF cells from OS samples represented in a t-SNE plot graph. Different cell type clusters were color coded. (B) Violin plots of selected apCAF and mCAF markers, showing normalized expression in two of the subclusters. (C) Many specific genes were uniquely upregulated in the apCAF subtype, including HLA-DRA, CD74, APOE *etc*.

### Functional analysis of the myeloid cells in OS

Tumor□infiltrating myeloid cells are the most abundant monocyte population within tumors and known for their functional and molecular plasticity. In this paper, we identified 7 subgroups of the myeloid cells, including the M1-like macrophage, M2-like macrophage, IFN activated macrophage, CD14+ monocyte, DC, proliferating myeloid cells and neutrophil cells (Fig. 5A). Each subgroup of the myeloid cells had distinctive biomarkers, which were shown in violin-plot (Fig. 5 B).

**Fig. 5.**
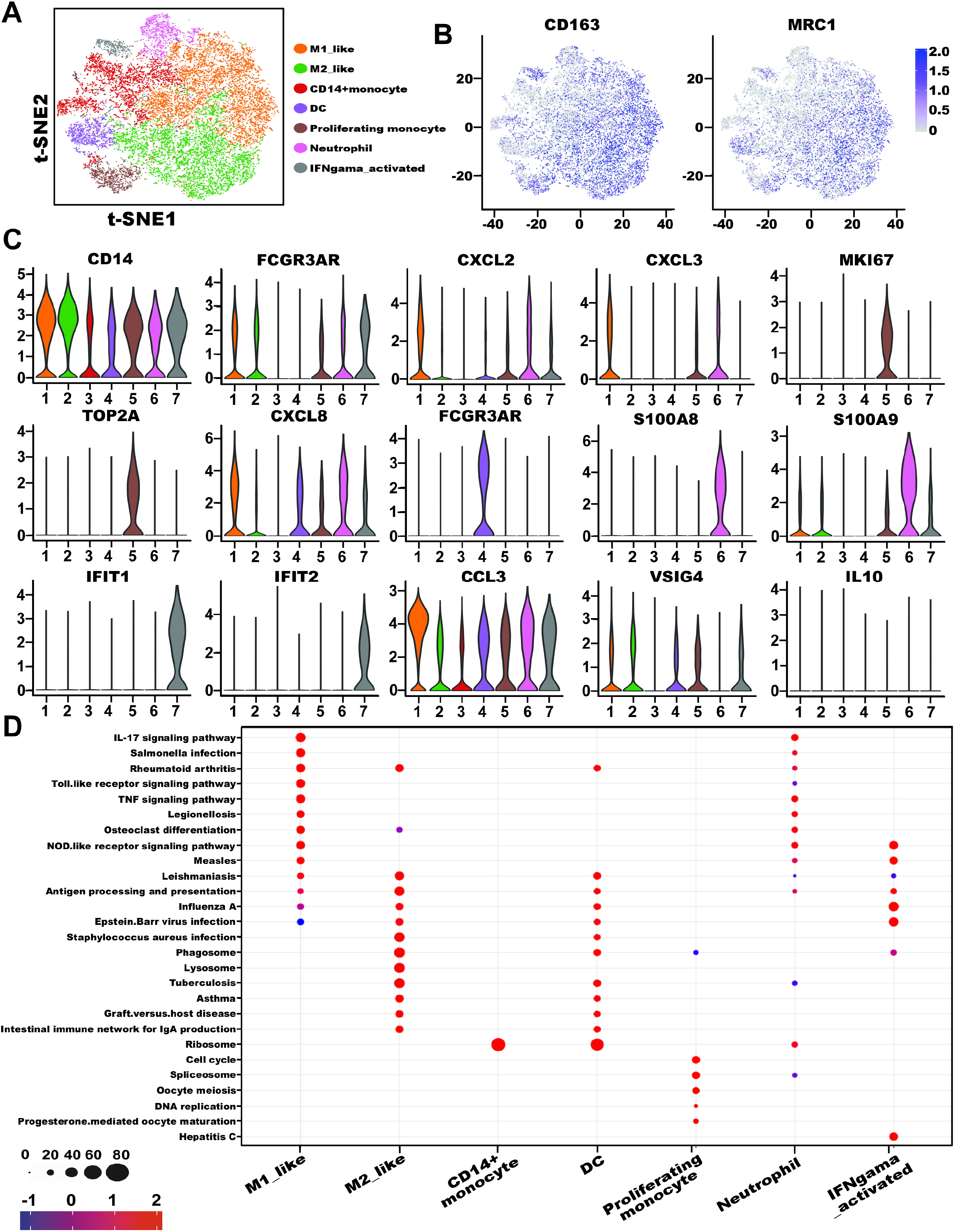
Single-cell analysis of myloid cells in OS samples. (A) t-SNE plot of the myeloid cell subgroup in OS tumor samples. (B) M2-polorized markers CD163 and MRC1/CD206 were expressed in almost all TAMs. (C) The relative expression levels of the well-known biomarkers of each cell type as indicated in the violin plots. (D) Bubble plot was used to identify each cell type–specific markers across clusters. Size of dots represents fraction of cells expressing a particular marker, and intensity of color indicates level of mean expression.

Tumor-associated macrophages (TAMs) are the major immune component of myeloid cells in OS. The majority of the TAMs have relatively higher expression level of CD163 and mannose scavenger receptor(MRC1)/CD206, suggesting that these cells were M2-polorized TAMs in OS (Fig. 5 C). The TAMs were clustered into 3 subgroups. The first group was the M1-like TAMs expressing a relatively higher level of pro-inflammatory markers including C-C motif chemokine ligand 2(CCL2), CCL3, CCL4, CXC motif chemokine ligand 2(CXCL2) and CXCL3. The main ingredient was M2-like TAMs with relatively higher expression of inflammatory biomarkers including IL-10. We also found the IFN activated macrophage, which was characterized with higher expression levels of IFN-induced proteins with tetratricopeptide repeat 1(IFIT1), IFIT2 and IFIT3, suggesting that the activation of IFN signaling pathway may contribute to the tumor suppressive microenvironment (Fig. 5D). Our data imply OS is abundant in M2-like immunosuppressive TAMs.

### Contribution of the immune-suppressive tumor-infiltrating Lymphocyte(TIL) cells to OS

The presence and content of TILs is considered to be closely related with response to the immunotherapy[25]. Here we characterized the subpopulations of TIL using transcriptomic patterns. According to the analysis, the major groups of the lymphoid cells included CD4-/CD8-T cells, CD8+T cells, NK cells, T-reg cells, mast cells and plasma cells (Fig.6A). The lymphoid cells were the majority in the primary tumor and the lung metastatic tumor. In contrast, they were rarely noted in the recurrence tumor samples. Most of the T cells in the primary OS tumor were CD4-/CD8-T and Treg cells. In the lung metastasis, about 311 out of the 1969 T cells were T-reg cells. Meanwhile, the cellular distribution of NK cells and the CD8+T cells were significantly reduced in the lung metastatic tumors. Using the IHC method (Fig. 6B), we validated the cellular composition of the lymphoid cells, and found that cytotoxic CD8+T cells barely existed in the recurrent and lung metastatic tumors. Our results suggested the cytotoxity of T cells is loss-of-function in the OS tumor, especially in recurrent and metastatic tissue.

**Fig. 6.**
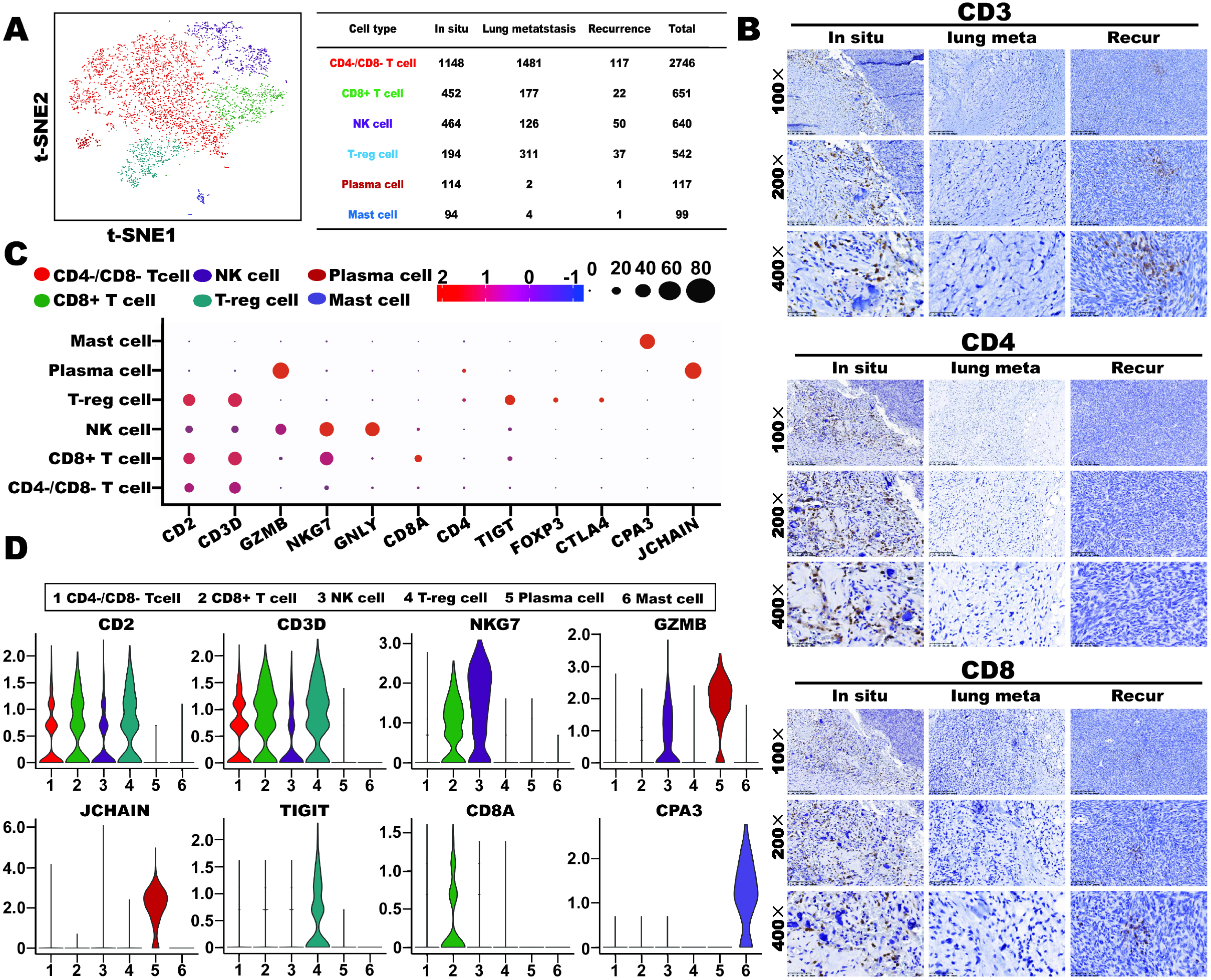
Subgroups of TILs in the OS tumor. (A) Reclustering of the subgroup of TILs in the OS data represented in the t-SNE plot. Proportion of each subgroup cells in primary, recurrent and pulmonary metastatic tumor samples was provided. The tab exhibited the concrete value of each scRNA-seq sample gained. (B) Microscopy of the expression of CD3, CD4 and CD8 in primary, recurrent and pulmonary metastatic OS tissues. Black scale bar is 100 μm. (C) The violin plots expound normalized levels of markers in the different subclusters. (D) The well-known markers of each cell population were represented in dot plot.

We performed the dot-plot analysis to display the level of marker genes in all kinds of cells (Fig. 6C). The GZMB expression level was reduced, suggesting that CD8+T cells in the OS cells had lower cytotoxic activities. The cellular composition of the T-reg cells was significantly increased, and the T-reg cells expressed relatively higher levels of cytotoxic T-lymphocyte-associated protein 4(CTLA-4) and TIGIT, which are negative regulator for the cytotoxicity of the CD8+T cells (Fig. 6D).

### Blocking TIGIT improved the cytotoxicity of Cytokine induced T cells (CITs) to OS cells

TIGIT is expressed normally by activated T cells, regulatory T cells (Treg), and natural killer (NK) cells, which is recently emerging as novel candidate in immunotherapy. As mentioned above, we show for the first time that the TIGIT over-expressed on TILs of OS patients by scRNA-seq. It was verified by IHC method(Fig. 7A). In order to demonstrate the therapeutic potential of TIGIT in OS, We isolated the CD3+ T cell and blocked the inhibitory activity of TIGIT. The immune cell-mediated lysis of CITs on OS cells was measurably enhanced by the addition of blocking TIGIT antibodies in co-culture system(*p*<0.05, Fig. 7B).

**Fig. 7.**
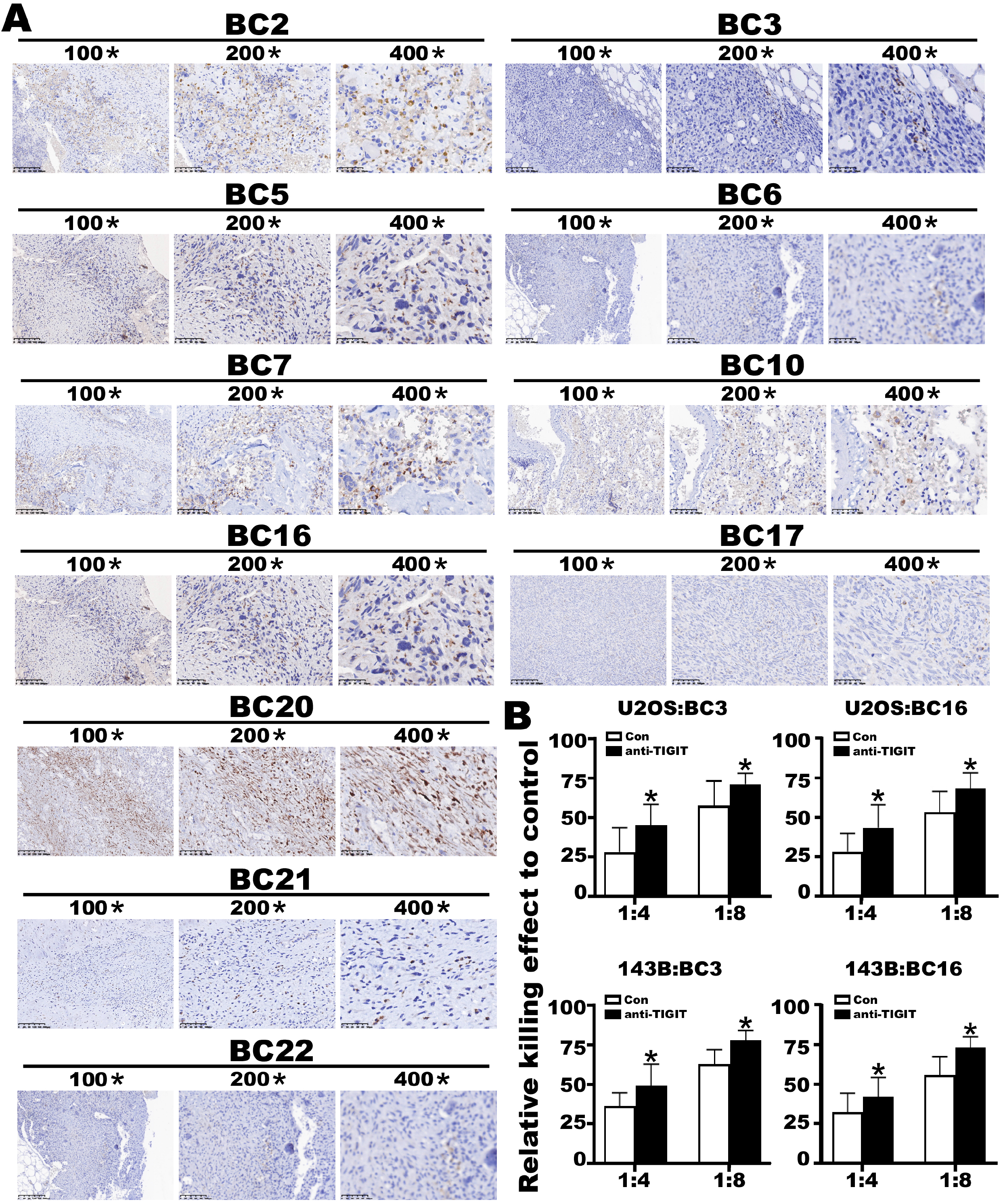
Blockade of the TIGIT increases the specific lysis of breast cancer cell lines. (A) Immunostaining of TIGIT in 11 OS tissue displayed dark brown. Scale bar: 100 μm. (B) We analyzed the lysis of the cytokine-induced killer cells (CIKs) produced from OS patients(n = 2) with or without blocking TIGIT on the OS cell line U2OS and 143B. Results of cytotoxicity ratio are depicted as the mean ± SD. For statistical analysis paired T-tests were performed (* means *p* < 0.05)

## Discussion

WES/WGS data or transcriptomic results had described OS is a highly inter-heterogeneous tumor[26]. However, they are only reflecting the average of expression levels across the tumor cells. As such, these studies could not identify cell types, nor predict developmental trajectories, clarify taxonomic composition and metabolic capacities of TME. In the present study, we applied scRNA-seq strategy to profile malignant cells and TME components from primary, recurrent and lung metastatic OS tissues. To our knowledge, this paper is the first study which performed scRNA-seq to identify the intra-heterogeneous of OS. Thus, how to identify the OS cells is most important. Han et al proposed that Ctsk+ cells serve as a physiologic and pathological precursor in osteogenic tumor[27–29], and we chose it as one of the major indicators. Meanwhile, bone sialoprotein (BSP) is thought to function in the initial mineralization of bone and selectively expresses by differentiated osteoblast[30]. Here, the OS cells were characterized using CDH11, BSP, LUM, DCN and COL1A1 as markers.

In the subgroup analysis, we put emphasis on the subtype 5 enriching in multiple singling pathways related to cancer development and other transcriptional misregulation signaling pathways. Meanwhile, the genes related to p53 downstream signaling pathways were significantly enriched in the subcluster. As we know, TP53 and Rb1 exhibited the most frequent mutations in OS, which may be the initiating factors for OS tumorigenesis[30]. The OS cells with TP53 mutation exhibit a high chromosomic instability and lead to secondary genetic aberrations in new cancer cell clones that emerged from the initial monoclone [31]. A large number of animal models are developed for osteosarcoma, including P53 knock out mouse model [32]. Based on our cellular trajectory analysis, the precursor of OS is highly proliferative. It is widely accepted that the stem cell could renew but being at G0 stage with proliferative inactivity. We were unable to capture the stem cells, probably due to their extremely low number. In our opinion, the differentiation subtype may be the secondary genetic aberrations, which would then transform to special metabolism, bone matrix remodeling and ossification subtypes. We hypothesize that the components in our paper behave just like myeloblast, promyelocyte and so on, thus presenting the hematopoietic cell differentiation and development processes. Of course more research is necessary to further explore this hyopthesis.

Emerging immune checkpoint inhibitors have been the landmark treatments for their clinical success in a variety of human cancers. However, the therapeutic efficacies by immune checkpoint inhibitors were variable for the treatment of OS patients[33, 34]. Gomez-Brouchet et al. reported that PD1/PDL-1 staining was negative in >80% of OS cases (n=124) [35]. Alves showed that OSJ displayed a microenvironment with low tumor infiltrating lymphocytes (TILs), and few cells exhibited immunotherapeutic targets CTLA-4 and PD-1[36]. Palmerini examined the TAMs by primary OS tissue microarray to evaluate the status of the immune-infiltrates in OS. Most cases presented TILs, which contained CD3+ (90%) and CD8+ (86%). Meanwhile, PD-L1 expression was found in 14% patients in immune-cells and 0% in tumoral cells [37]. Our results were consistent with theirs: low cytotoxic TILs were usually in OS and the level of PD-L1 was so low that it could not be detected in OS tissue. On the contrary, some previous studies on OS samples reported a higher rate of positive expression of PD-L1 (IC), ranging from 25% to 74% [38, 39]. We reason that these variant conclusions could be ascribed to insufficient statistical samples and different agents. In order to comprehensively predict the validity of ICIs, more indicators are required.

Wang reported that metastatic OS tumors showed improved immunogenicity. But most of TILs in lung metastasis were the naïve T cells or T-regs with lower anti-cancer activities[40]. These results thus appeared inconsistent. In our research, we found that the immunosuppression of lung metastasis tissue was more significant with the higher percentage of T-reg cells. In fact, it is controversial whether metastatic OS tumor was immunogenic. Our view is that significant immunophenotypic disturbances were found in recurrent and metastatic tissues, with immune-cells being rarely noted in recurrence OS tissue. Our data suggested that the OS immune environment became “cool” in the recurrent OS tumor. Thus, we initiated the clinical trial NCT03676985 using anti-PD-L1 as maintenance after adjuvant chemotherapy.

Another key finding of our paper is that the T-reg cells had a relatively higher TIGIT expression level in OS tumor. TIGIT and its ligand poliovirus receptor (PVR) have been emerging as novel promising targets in immunotherapy for many tumors such as breast cancer, lung cancer, hepatocellular carcinoma etc[41–43]. Tian reported that blockade of TIGIT prevented NK cell exhaustion and elicited potent anti-tumor immunity[44]. In the current study, we observed that the T-reg cells had relatively higher TIGIT expression level in OS tumor and blockade of TIGIT improved the cytotoxicity of CIT, which suggested that OS patients may benefit from individualized immunotherapy according to the genetic results.

TAMs are known to participate in tumor initiation, progression and metastasis. We found that the majority of the TAMs have relatively higher CD206 and MRC1 expression, suggesting that TAMs were M2-type with anti-inflammatory activities in OS tumor. Interestingly, we also observed a subgroup of TAMs with relatively higher expression level of inflammatory factors including CCL3, CCL4, CXCL8 etc., suggesting that the inflammatory activities may be involved in the tumorigenesis and progression of OS tumor. Our scRNA-seq analysis displayed the dynamic development of TAMs, which was consistent with previous studies reported by Dumars et al. that M2-macrophages were dominated in OS tissue and that macrophage dyspolarization was associated with metastatic process in OS patients [45]. In this regard, our data seems to offer some hints as to why the mifamurtide, a fully synthetic lipophilic derivative of the muramyl dipeptide (MDP) encapsulated into liposomes, was effective when used together with chemotherapy for localized OS; hence, the addition of mifamurtide was preferred in non-metastatic OS patients, whereas there was no significant difference in overall survival rates between the combined use of mifamurtide with chemotherapy and chemotherapy alone in metastatic OS [46].

In recent years, CAFs have attracted attention due to their role in mediating collagen crosslinking with malignant cells by disintegrating metalloproteinases (ADAMs) and secreting multiple cytokines, chemokines and growth factors[47]. Meanwhile, CAFs assist cancer cells in evading immune surveillance through inhibiting the activity of immune-effector cells and recruiting immune-suppressive cells, thus supporting cancer tumorigenesis and metastasis[48]. However, the role of the CAFs in driving tumorigenesis of OS remains to be further elucidated. As mentioned above, Elyada et al[49] described a new population of CAFs that displayed relatively higher expression of MHC class II and CD74 without the expression of classic costimulatory molecules, and they named it “antigen-presenting CAFs”. In our study, we also isolated two subtypes of CAFs, apCAFs and the traditional myCAFs. Previous studies suggested that apCAFs could activate CD4+ T cells and act as the immune-modulator [50]. We found the ratio of apCAF/myCAF in primary tumor tissue was dramatically higher than that in pulmonary metastatic and recurrent tumor tissues, suggestive of the difference of the TME of OS in the primary *versus* the metastatic tumor. Nonetheless, more studies are warranted to address the origin of the apCAFs and their roles in the OS progression and development.

In summary, using the scRNA-seq method, our study uncovers the intra-heterogeneity of both malignant OS cells and the TME. Distinct subgroups of OS cells were documented and the cellular lineage in lung metastasis of OS were determined. Furthermore, the main immune cell types in the TME were profiled. Together, our findings in the present study may provide novel therapeutic targets and methods for the treatment of OS patients.

## ACKNOWLEDGMENTS

We thank all the patients who contributed to this study. We also thanks the Hengrui pharmaceutical co. LTD due to providing the antibodys of TIGIT and CTSK for free.

---

**Fig. S1. The t-SNE analysis of individual**

(A) The t-SNE results of 11 samples were shown on the list. The cell number and percentage of assigned cell types were summarized in the right panel. Cell number of all clusters were summarized in the right tab. (B) The t-SNE were gathered by region.

**Fig. S2. The analysis of key genes in OS cells**

(A) The KEGG analysis of OS cells was shown on the list. (B) The remarkable genes of each subtype were enriched in heatmep.

**Fig. S3. The CNV analysis of OS cells**

(A) The CNV analysis of primary OS cells revealed amplication of the chromosome 21 and 7 in almost all samples. (B) The CNV analysis of lung metastatic OS cells showed rare del in all chromosomes. (C) The CNV analysis of recurrent OS cells also exhibited amplication of the chromosome 21,7.

**Fig. S4. The t-SNE analysis of TILs from OS different location**

(A) The t-SNE results of TILs from primary OS tissue. (B) The t-SNE results of TILs from lung metastasis OS tissue. (C) The t-SNE results of TILs from recurrent OS tissue.

**Fig. S5. The co-expression of CD74 and CTSK in OCs at different OS tissue**

(A) The OC cells with CD74 and CTSK expression in primary OS tissue. (B) The OC cells with CD74 and CTSK expression in recurrent OS tissue. (C) The OC cells with CD74 and CTSK expression in lung metastasis OS tissue.(The CD74 positive cell presented red color, the CTSK positive cell presented green color, the nuclear was stain blue color)

