## Supplemental1 for "Single-cell RNA-seq reveals identity and heterogeneity of malignant osteoblast cells and TME in osteosarcoma"

### Figure S1

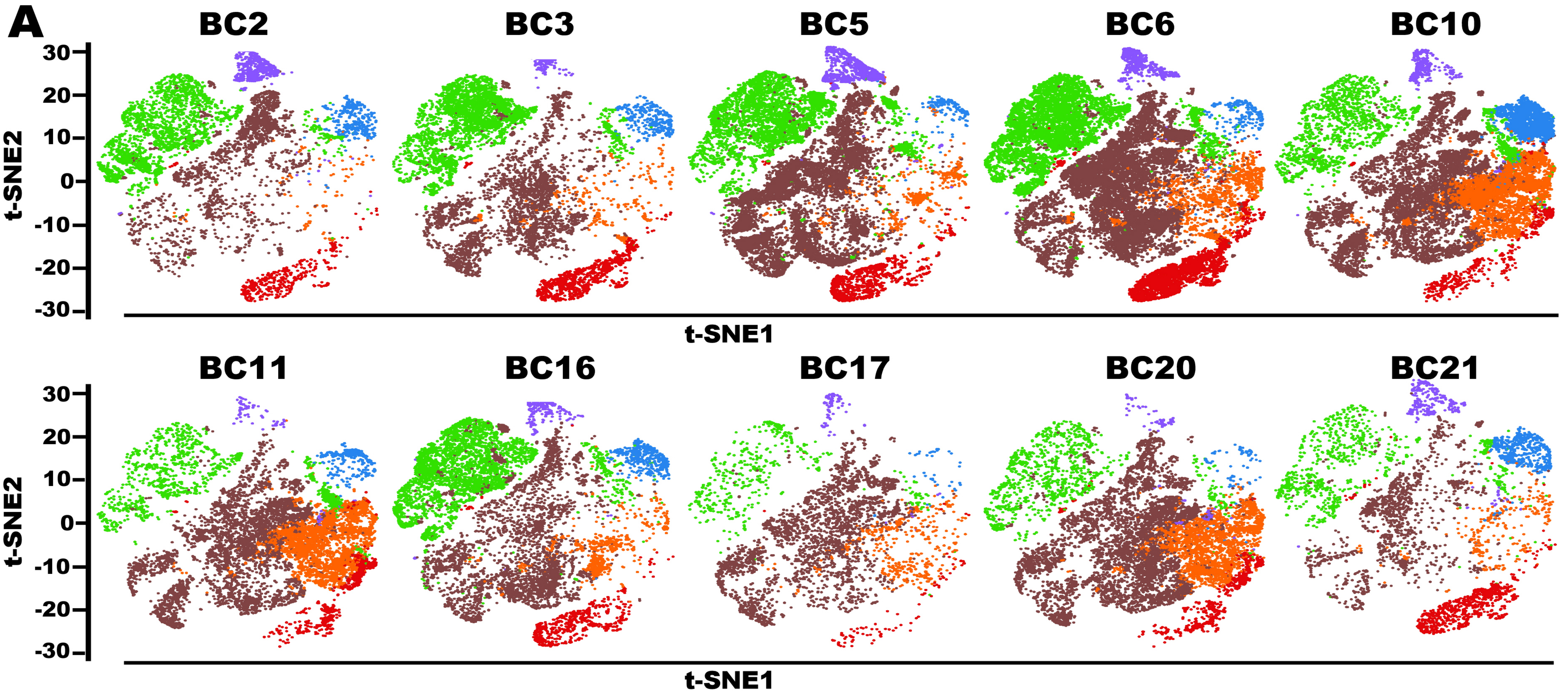


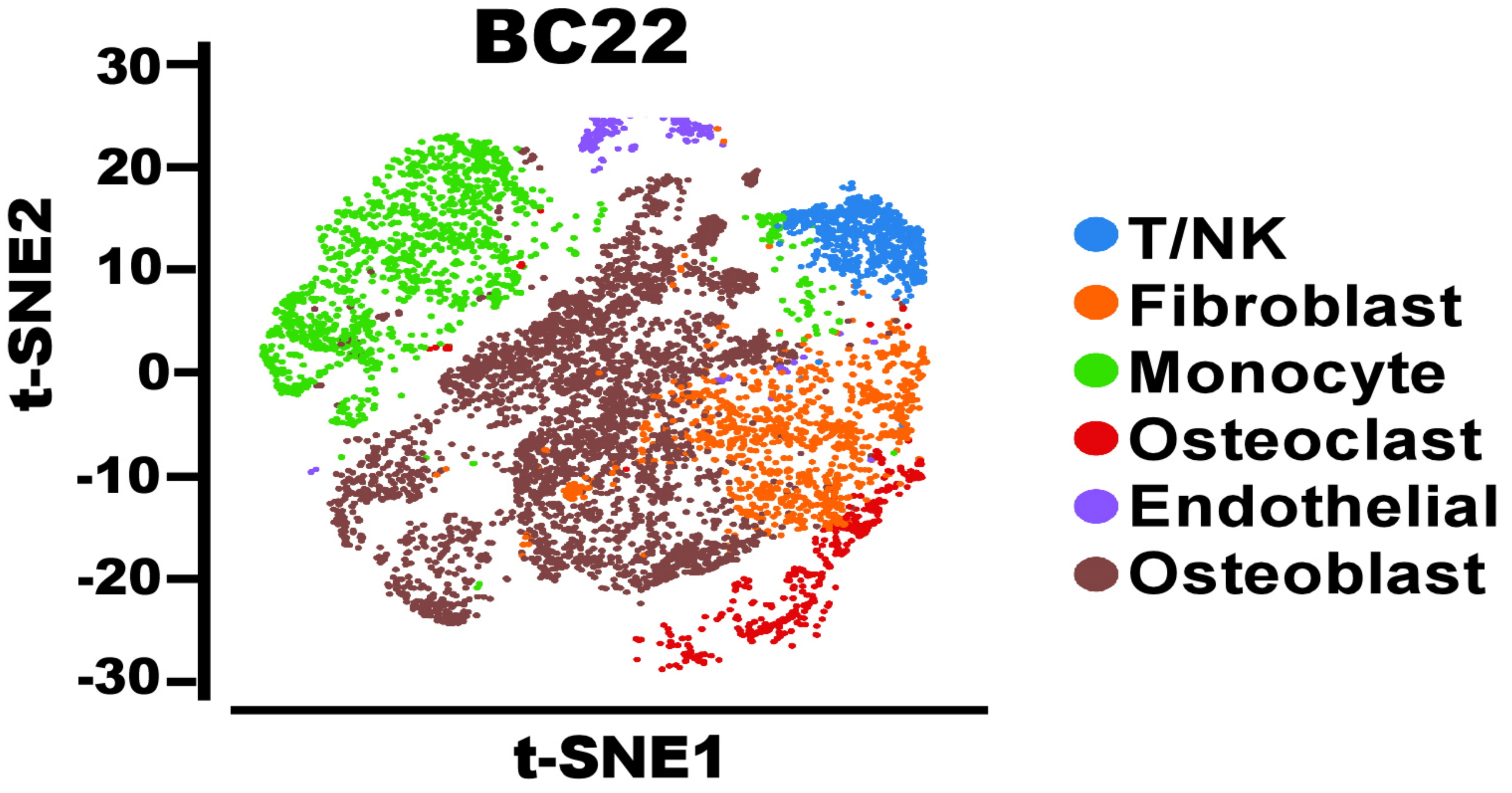


#### Number of cell

| Cell Type | Insitu | Lung metastasis | Recurrence | Total |
|---|---|---|---|---|
| T/NK_cell | 2941 | 2139 | 336 | 5416 |
| Fibroblast_cell | 7180 | 5262 | 5788 | 18158 |
| Monocyte | 22033 | 2438 | 2301 | 26772 |
| Osteoclast | 7755 | 640 | 785 | 9180 |
| Endothelial | 7755 | 640 | 785 | 9180 |
| Osteoblast cell | 29230 | 8520 | 9848 | 47598 |

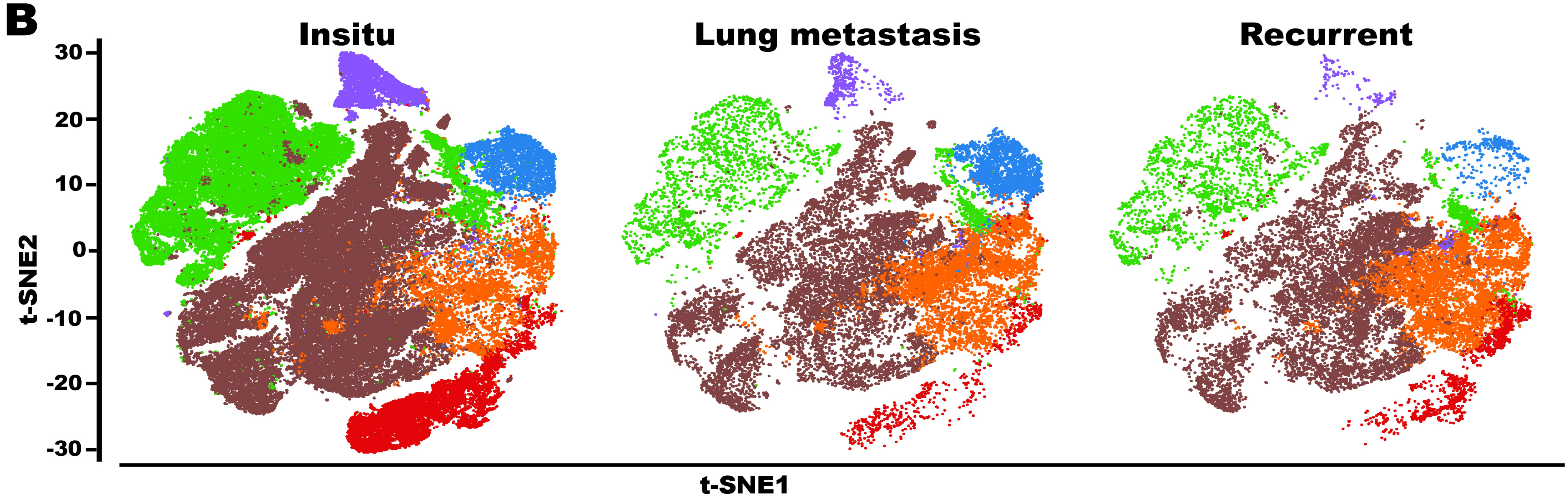
