## Supplementary figures and images for "Single-cell RNA-seq reveals identity and heterogeneity of malignant osteoblast cells and TME in osteosarcoma"

### Supplemental2

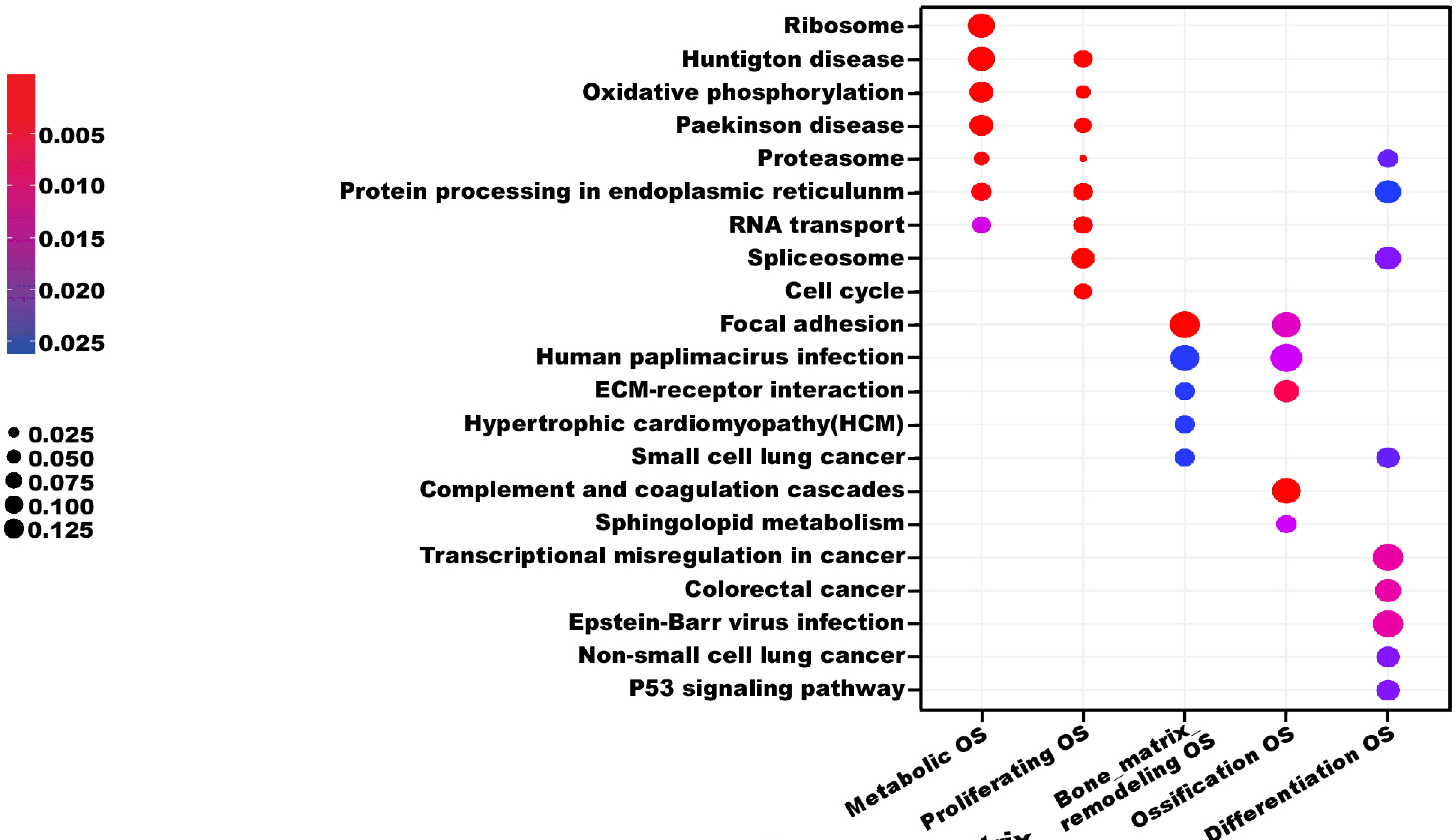

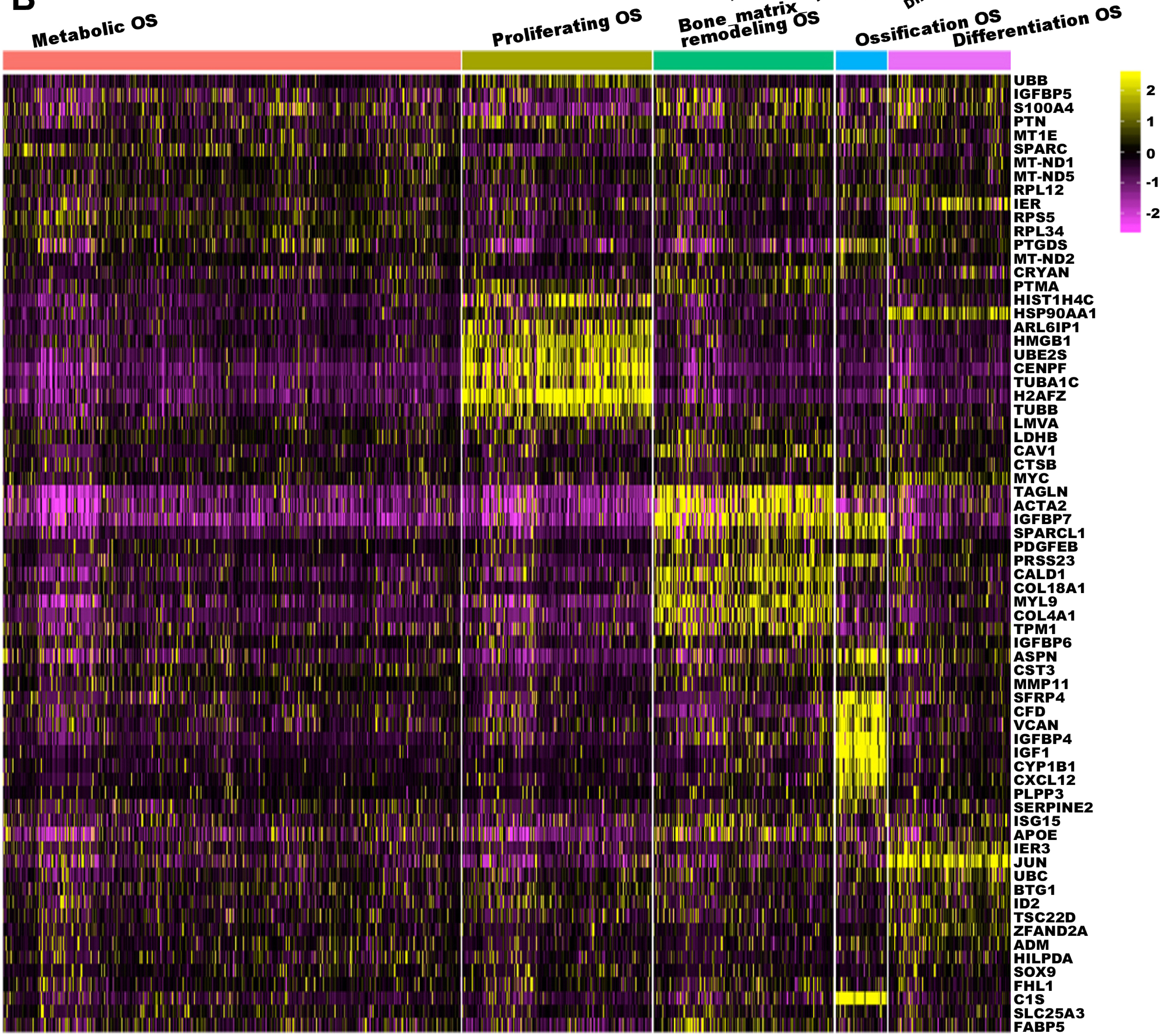

### Supplemental3

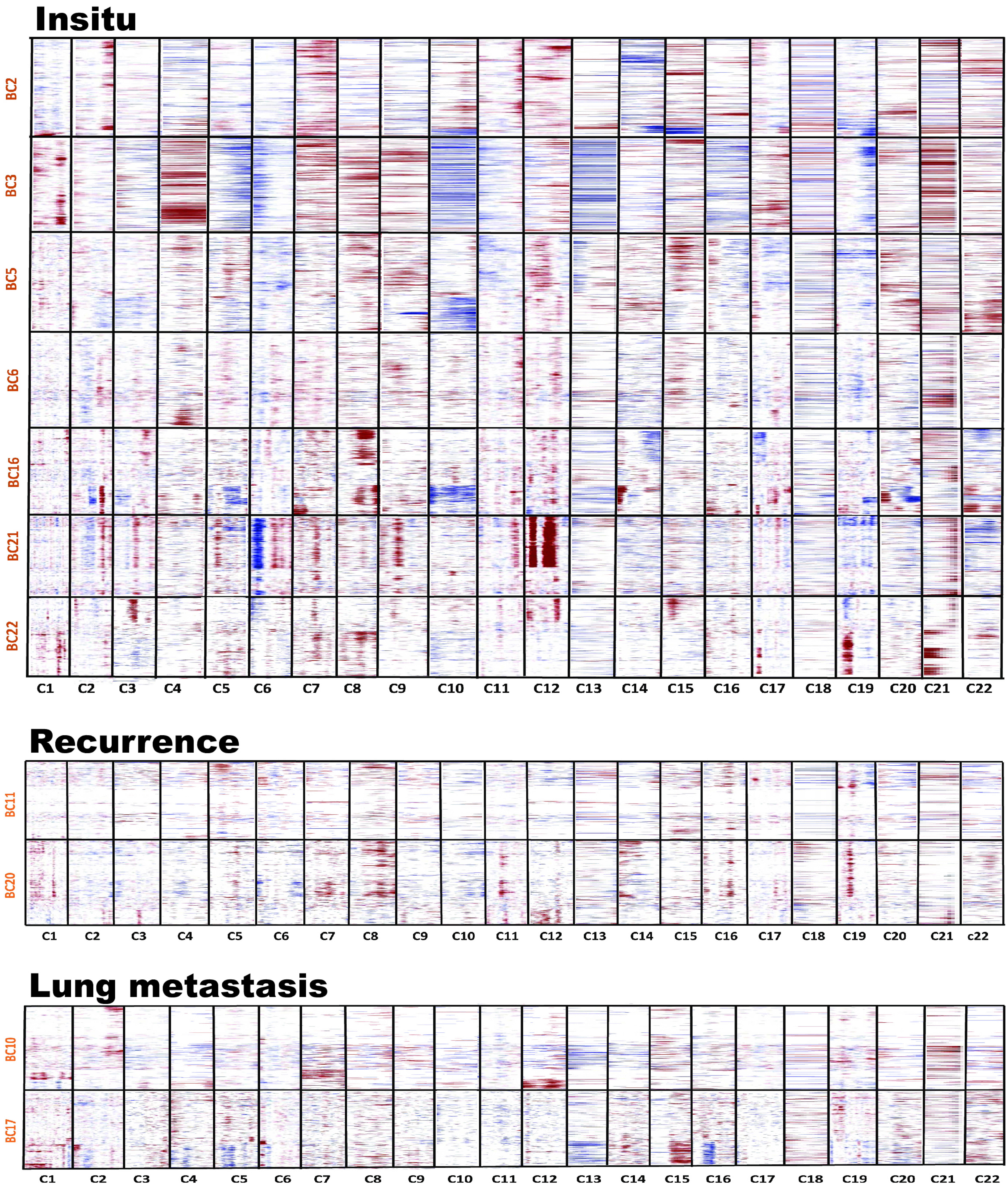

### Supplemental4

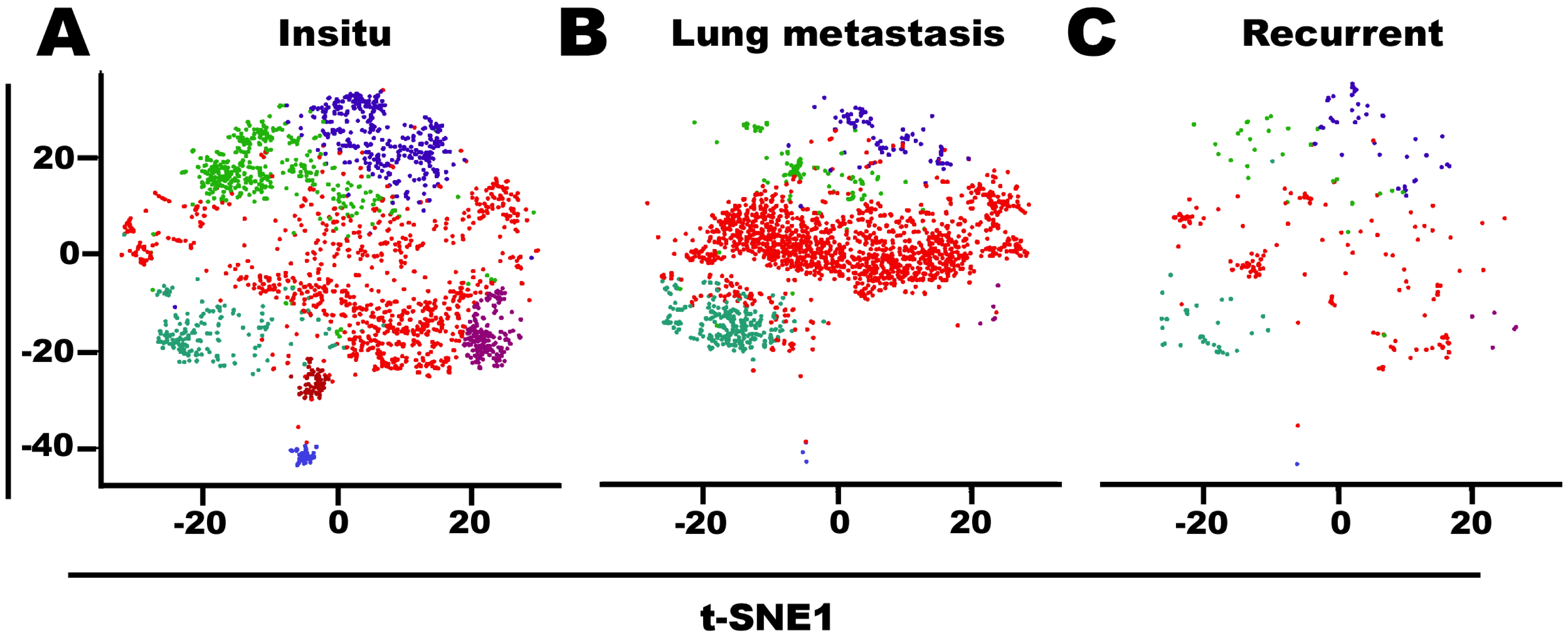

### Supplemental5

#### Figure S5

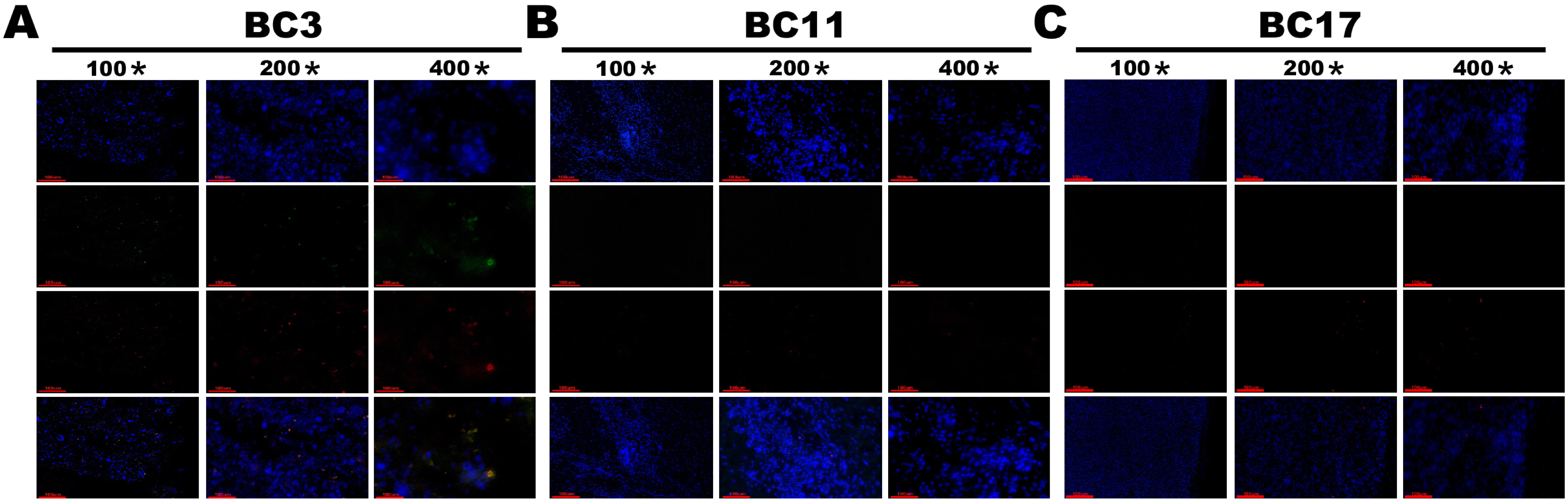
